# Bacterial profiles of the human placenta from term and preterm deliveries

**DOI:** 10.1101/2022.06.21.497119

**Authors:** Kevin R. Theis, Andrew D. Winters, Roberto Romero, Ali Alhousseini, Jonathan M. Greenberg, Jonathan Panzer, Jose Galaz, Percy Pacora, Zachary Shaffer, Eunjung Jung, Nardhy Gomez-Lopez

## Abstract

Whether the human placenta is a sterile organ is under debate. Yet, infection of the amniotic cavity, including the placenta, is causally linked to preterm birth. This study compares the bacterial profiles of term and preterm placentas through culture and 16S rRNA gene sequencing of the amnion, amnion-chorion interface, subchorion, villous tree, and basal plate, while accounting for patient identity, mode of delivery, presence/absence of labor, and potential background DNA contamination. As no evidence of a placental microbiota in term pregnancy was found, these placentas were considered as controls. Placentas from preterm birth cases were more likely to yield bacterial cultures, and their bacterial DNA profiles were less rich than those of term controls, suggesting the predominance of only a few bacteria. Nevertheless, the bacterial DNA profiles of placentas from preterm cases and term controls were not consistently different. The placentas from preterm cases may often have a microbiota but the bacteria constituting these communities varied among the women. Mode of delivery had a pronounced effect on the bacterial profiles of all sampled levels of the placenta. Specifically, the bacterial DNA profiles of vaginally delivered placentas had higher relative abundances of *Finegoldia*, *Gardnerella*, *Peptoniphilus*, and *Prevotella* (each a common resident of the vaginal microbiota) than the profiles of cesarean-delivered placentas. Collectively, these data indicate that there is a not a placental microbiota in normal term pregnancy, and that although the placentas of some preterm cases were populated by bacteria, the identities of these bacteria varied among women delivering preterm.

**IMPORTANCE:** If a placental microbiota exists, then current understanding of the roles of microorganisms in pregnancy outcomes need to be reconsidered. For instance, we will need to determine if a placental microbiota is beneficial to pregnancy outcome by excluding potential pathogens from colonizing the placenta and/or effectively priming the fetal immune system, and furthermore which characteristics of the placental microbiota preclude versus promote placental infection, which can result in pregnancy complications such as preterm birth. Our findings here are consistent with prior investigations that have reported that there is not a placental microbiota in typical human pregnancies. Yet, bacteria can be detected in placentas from preterm deliveries. The principal source of microorganisms invading the amniotic cavity, including the placenta, is the vaginal microbiota. Focus should be on elucidating the metabolic and/or virulence characteristics of the subset of bacteria within the vaginal microbiota that commonly invade the amniotic cavity, resulting in infection.

## INTRODUCTION

Whether there exists a low biomass microbiota (i.e., resident bacterial community) in human placentas from uncomplicated pregnancies is under debate (1–14). The placenta has typically been viewed as a sterile organ (5). Yet, it has long been known that bacterial infection of the placental amnion and chorion (15–24) and/or villous tree (15, 18, 25–27) is associated with preterm labor (16, 18, 28–34), preterm prelabor rupture of the membranes (PPROM) (18, 29, 30, 35, 36), histological chorioamnionitis (15-17, 22, 23, 37, 38), clinical chorioamnionitis (15, 36–42), and congenital infection (26, 43–47). What is unique about many recent investigations is that they further report detection of a microbiota in placentas from uncomplicated pregnancies at term (1, 2, 4, 10, 48–58), and have concluded that the bacterial profiles of placentas from pregnancies complicated by spontaneous preterm birth (1, 48, 52), severe chorioamnionitis (52, 53), gestational diabetes mellitus (51), and low (50) or high (55) neonatal birthweight differ from the bacterial profiles of placentas from uncomplicated pregnancies at term. If true, this would be a paradigm shift – all human placentas have a resident microbiota; however, the structure of the microbiota varies with different pregnancy complications.

The verified existence of a placental microbiota absent infection would require a fundamental reconsideration of the roles of microorganisms in human pregnancy outcomes. Most importantly, it would need to be determined which characteristics of a placental microbiota (e.g. taxonomic composition and/or absolute abundance) preclude versus promote placental infection and pregnancy complications (48, 49). Additionally, it would need to be determined if a placental microbiota is typically inconsequential to pregnancy outcome or if it is potentially beneficial to its human host by competitively excluding placental colonization by pathogens and/or priming the fetal immune system for the microbial bombardment to be experienced upon delivery.

However, the existence of a placental microbiota is controversial (5, 6, 8, 13, 14). The crux of the debate is that most of the recent investigations proposing the existence of a placental microbiota have relied exclusively on DNA sequencing to detect and characterize bacterial communities in the placenta (1, 4, 48–58), and typically have not accounted for the potential influence of background DNA contamination on sequencing results (6, 59–62). Indeed, recent studies that have accounted for background DNA contamination have not found evidence of a placental microbiota in uncomplicated pregnancies at term (3, 7, 9, 11, 63–65). Instead, they support the classical paradigm – the human placenta is typically sterile (5), yet it can become infected and this can result in pregnancy complications such as preterm birth (15, 16, 18, 28–36), the leading cause of neonatal morbidity and mortality worldwide (66–68).

In this study, we compare and contrast the bacterial profiles of placentas from term and preterm deliveries. Specifically, we cultured bacteria from the placentas of term and preterm deliveries, and characterized the bacterial profiles of the amnion, amnion-chorion interface, subchorion, villous tree, and basal plate from term and preterm deliveries using 16S rRNA gene sequencing, accounting for individual identity, mode of delivery and the presence/absence of labor. As in prior investigations by our group and others (3, 7, 9, 11, 63–65), we found no evidence of a placental microbiota in uncomplicated pregnancies at term. Viable bacteria were recovered in culture mostly from term and preterm placentas that were vaginally delivered and, to a lesser extent, from preterm placentas obtained via a cesarean section. Yet, there was no consistent effect of gestational age at delivery on the overall 16S rRNA gene profiles of the amnion, amnion-chorion interface, subchorion, villous tree, and basal plate. Notably, various bacteria were identified in placentas from preterm cesarean deliveries through culture and molecular microbiological techniques. Since these placentas were not subject to potential influences of bacterial contamination from the vaginal delivery process, these findings suggest a bacterial presence in placentas in some cases of preterm birth.

## RESULTS

### Demographics

This prospective cross-sectional study included 69 patients, 49 of whom delivered at term and 20 of whom delivered preterm. Term deliveries included 20 cesarean not in labor (NIL), 8 cesarean in labor (IL), and 21 vaginal deliveries. Preterm deliveries included 9 cesarean NIL, 5 cesarean IL, and 6 vaginal deliveries. Patient demographic and clinical characteristics are summarized in **Table 1.**

**Table 1.**
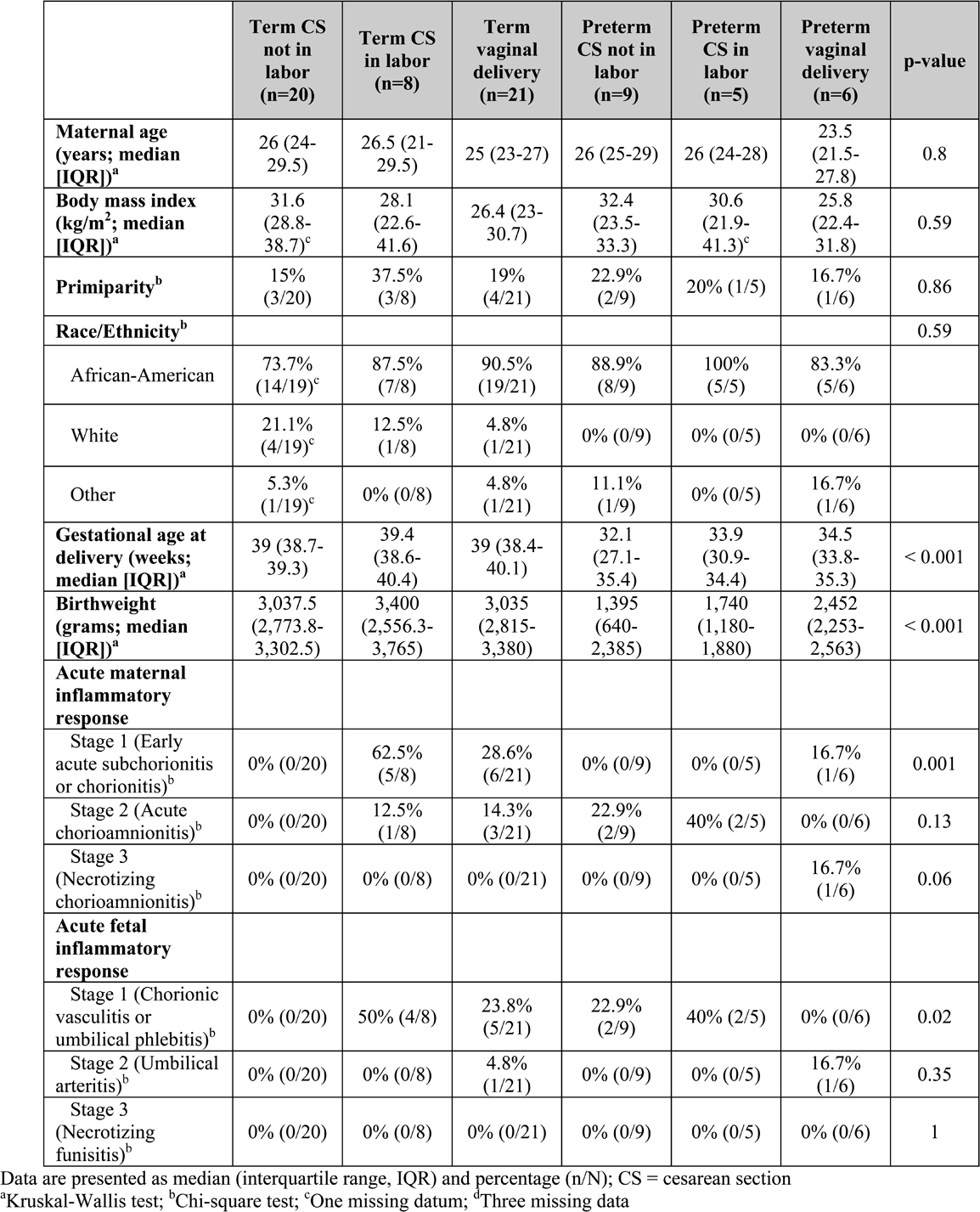
Demographic and clinical characteristics according to the mode of delivery.

|  | Term CS<br>not in<br>labor<br>(n=20) | Term CS<br>in labor<br>(n=8) | Term<br>vaginal<br>delivery<br>(n=21) | Preterm<br>CS not in<br>labor<br>(n=9) | Preterm<br>CS in<br>labor<br>(n=5) | Preterm<br>vaginal<br>delivery<br>(n=6) | p-value |
| --- | --- | --- | --- | --- | --- | --- | --- |
| **Maternal age**<br>**(years; median**<br>**[IQR])<sup>a</sup>** | 26 (24-29.5) | 26.5 (21-29.5) | 25 (23-27) | 26 (25-29) | 26 (24-28) | 23.5<br>(21.5-27.8) | 0.8 |
| **Body mass index**<br>**(kg/m<sup>2</sup>; median**<br>**[IQR])<sup>a</sup>** | 31.6<br>(28.8-38.7)<sup>c</sup> | 28.1<br>(22.6-41.6) | 26.4 (23-30.7) | 32.4<br>(23.5-33.3) | 30.6<br>(21.9-41.3)<sup>c</sup> | 25.8<br>(22.4-31.8) | 0.59 |
| **Primiparity<sup>b</sup>** | 15%<br>(3/20) | 37.5%<br>(3/8) | 19%<br>(4/21) | 22.9%<br>(2/9) | 20% (1/5) | 16.7%<br>(1/6) | 0.86 |
| **Race/Ethnicity<sup>b</sup>** |  |  |  |  |  |  | 0.59 |
| African-American | 73.7%<br>(14/19)<sup>c</sup> | 87.5%<br>(7/8) | 90.5%<br>(19/21) | 88.9%<br>(8/9) | 100%<br>(5/5) | 83.3%<br>(5/6) |  |
| White | 21.1%<br>(4/19)<sup>c</sup> | 12.5%<br>(1/8) | 4.8%<br>(1/21) | 0% (0/9) | 0% (0/5) | 0% (0/6) |  |
| Other | 5.3%<br>(1/19)<sup>c</sup> | 0% (0/8) | 4.8%<br>(1/21) | 11.1%<br>(1/9) | 0% (0/5) | 16.7%<br>(1/6) |  |
| **Gestational age at**<br>**delivery (weeks;**<br>**median [IQR])<sup>a</sup>** | 39 (38.7-39.3) | 39.4<br>(38.6-40.4) | 39 (38.4-40.1) | 32.1<br>(27.1-35.4) | 33.9<br>(30.9-34.4) | 34.5<br>(33.8-35.3) | < 0.001 |
| **Birthweight**<br>**(grams; median**<br>**[IQR])<sup>a</sup>** | 3,037.5<br>(2,773.8-3,302.5) | 3,400<br>(2,556.3-3,765) | 3,035<br>(2,815-3,380) | 1,395<br>(640-2,385) | 1,740<br>(1,180-1,880) | 2,452<br>(2,253-2,563) | < 0.001 |
| **Acute maternal**<br>**inflammatory**<br>**response** |  |  |  |  |  |  |  |
| Stage 1 (Early<br>acute subchorionitis<br>or chorionitis)<sup>b</sup> | 0% (0/20) | 62.5%<br>(5/8) | 28.6%<br>(6/21) | 0% (0/9) | 0% (0/5) | 16.7%<br>(1/6) | 0.001 |
| Stage 2 (Acute<br>chorioamnionitis)<sup>b</sup> | 0% (0/20) | 12.5%<br>(1/8) | 14.3%<br>(3/21) | 22.9%<br>(2/9) | 40% (2/5) | 0% (0/6) | 0.13 |
| Stage 3<br>(Necrotizing<br>chorioamnionitis)<sup>b</sup> | 0% (0/20) | 0% (0/8) | 0% (0/21) | 0% (0/9) | 0% (0/5) | 16.7%<br>(1/6) | 0.06 |
| **Acute fetal**<br>**inflammatory**<br>**response** |  |  |  |  |  |  |  |
| Stage 1 (Chorionic<br>vasculitis or<br>umbilical phlebitis)<sup>b</sup> | 0% (0/20) | 50% (4/8) | 23.8%<br>(5/21) | 22.9%<br>(2/9) | 40% (2/5) | 0% (0/6) | 0.02 |
| Stage 2 (Umbilical<br>arteritis)<sup>b</sup> | 0% (0/20) | 0% (0/8) | 4.8%<br>(1/21) | 0% (0/9) | 0% (0/5) | 16.7%<br>(1/6) | 0.35 |
| Stage 3<br>(Necrotizing<br>funisitis)<sup>b</sup> | 0% (0/20) | 0% (0/8) | 0% (0/21) | 0% (0/9) | 0% (0/5) | 0% (0/6) | 1 |

Data are presented as median (interquartile range, IQR) and percentage (n/N); CS = cesarean section
<sup>a</sup>Kruskal-Wallis test; <sup>b</sup>Chi-square test; <sup>c</sup>One missing datum; <sup>d</sup>Three missing data

### Bacterial culture of placental tissues

The results of bacterial culture of placental tissues from the six patient groups are summarized in **Figure 1**. When a potential effect of labor was included in the multiple logistic regression analysis, mode of delivery and the interaction between mode of delivery and gestational age significantly affected the likelihood of obtaining a positive culture from placental tissues (mode of delivery: *p* = 0.002; mode of delivery*gestational age: *p* = 0.021). No placenta from a term cesarean delivery, regardless labor status, yielded even a single bacterial isolate **(Figure 1)**. In contrast, all three patient groups experiencing preterm deliveries (i.e., cesarean NIL, cesarean IL, and vaginal) included placentas that yielded bacterial isolates. This explains the significant interaction between mode of delivery and gestational age. *Escherichia coli* was recovered from placentas in each of the three preterm patient groups, yet it was not recovered from any placentas delivered at term **(Figure 1)**. When a potential effect of labor was removed from the multiple logistic regression analysis (due to consideration of a potential lack of statistical power), placentas from vaginal deliveries were more likely to yield a bacterial isolate than those from cesarean deliveries [odds ratio (OR) 7.65, 95% confidence interval (CI) 1.87-31.33, *p* < 0.001], and placentas from preterm deliveries were more likely to yield a bacterial isolate than those from term deliveries [(OR) 1.71, (CI) 0.48-6.05, *p* = 0.008].

**Figure 1.**
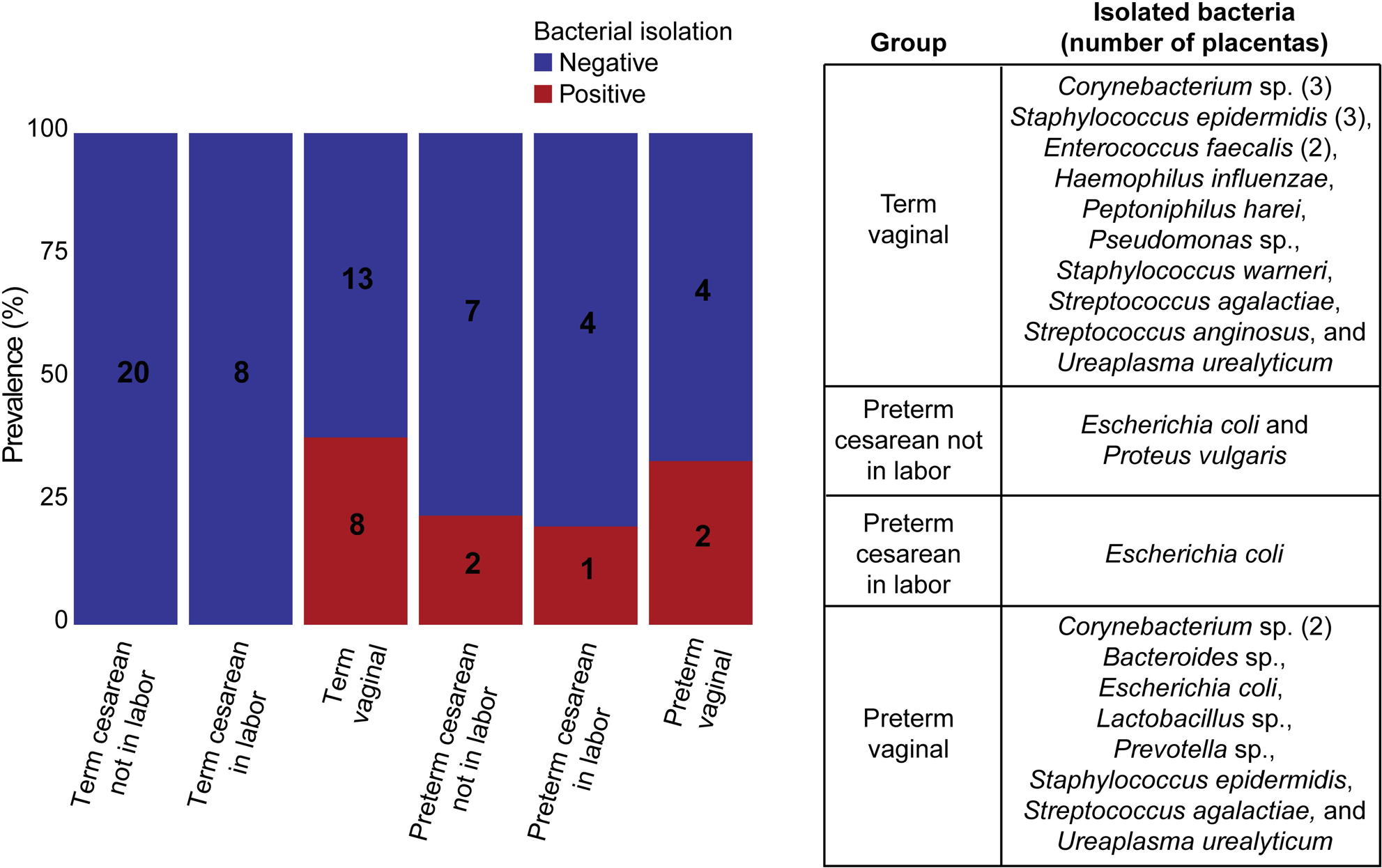
Results of bacterial culture of placental tissues from the different patient groups. The labels on bars indicate the number of patients within each group.

### Factors affecting the generation of a 16S rRNA gene sequence library at 35 cycles of PCR amplification

When a potential effect of labor was included in the multiple logistic regression model, no factors were identified as influencing the likelihood of generating a 16S rRNA gene sequence library from placental swabs at 35 cycles of PCR amplification (*p* ≥ 0.05). However, when a potential effect of labor was removed from the multiple logistic regression analysis, placental level (i.e., amnion, amnion-chorion interface, subchorion, villous tree, basal plate) was the lone factor influencing the likelihood of generating an amplicon library at 35 cycles (*X^2^* = 15.344, *df* = 4, *p* = 0.004). Pairwise comparisons **(Table 2)** revealed that swabs of the amnion were more likely to generate an amplicon library than swabs of the amnion-chorion interface, subchorion, or villous tree (*p* ≤ 0.003). Additionally, swabs of the basal plate were more likely to generate an amplicon library than were those of the subchorion or villous tree (*p* ≤ 0.001). However, swabs of the amnion were not more likely to generate an amplicon library than were those of the basal plate (*p* = 0.382). Collectively, these data indicate that the most exterior levels of the placenta (i.e., amnion, basal plate) – those levels that were most likely to be exposed to bacteria during delivery and/or sample processing – were most likely to generate an amplicon library at 35 cycles of PCR. Regardless, only samples that successfully amplified at 35 cycles of PCR amplification were included in subsequent analyses.

**Table 2.** Post-hoc analyses of the effect of placental level on the successful generation of 16S rRNA gene libraries at 35 cycles of PCR amplification. The analyses are Tukey’s pairwise comparisons with Bonferroni corrections applied for the model *16S rDNA library build success = MD*PL + MD*GA + PL*GA + (Patient ID)*, where *MD* is the mode of delivery, *PL* is the placental level, and *GA* is the gestational age at delivery. Patient ID was treated as a random variable.

| Linear hypothesis | Estimate | Standard error | *p*-value |
| --- | --- | --- | --- |
| Amnion-chorion interface vs. amnion | -1.992 | 0.547 | **0.003** |
| Basal plate vs. amnion | -1.163 | 0.561 | 0.382 |
| Basal plate vs. amnion-chorion interface | 0.830 | 0.425 | 0.508 |
| Subchorion vs. amnion | -2.785 | 0.531 | **< 0.001** |
| Subchorion vs. amnion-chorion interface | -0.793 | -2.195 | 0.281 |
| Subchorion vs. basal plate | -1.623 | 0.400 | **0.001** |
| Villous tree vs. amnion | -2.834 | 0.534 | **< 0.001** |
| Villous tree vs. amnion-chorion interface | -0.842 | 0.370 | 0.229 |
| Villous tree vs. basal plate | -1.671 | 0.406 | **< 0.001** |
| Villous tree vs. subchorion | -0.049 | 0.334 | 1.000 |

### Comparison of culture and 16S rRNA gene sequence data from placental samples

Strong concordance was observed between the bacterial culture and 16S rRNA gene sequence data of placental samples that yielded bacterial isolates **(Figure 2)**. For this analysis, placental swab sequencing libraries were included even if they had less than 500 sequences. Except for a single bacterial isolate (*Proteus vulgaris*), the bacteria (26/27; 96.3%) cultured from placental tissues had amplicon sequence variants (ASVs) with matching taxonomic classifications in at last one of the 16S rRNA gene profiles of their respective patient’s placental swabs. For the two preterm cesarean delivery cases in which *Escherichia coli* isolates were obtained, ASVs classified as *Escherichia* (99.6% shared nucleotide identity with *E. coli* via BLAST) had a relative abundance greater than 90% in at least four of the five sampled placental levels (i.e., amnion, amnion-chorion interface, subchorion, villous tree, basal plate). Other bacterial isolates that had high average relative abundances across their respective patient’s placental bacterial profiles were *Ureaplasma urealyticum* (62.7% and 43.5% in 2 patients), *Prevotella* sp. (33.5%), *Lactobacillus* sp. (13.4%), and *Haemophilus influenzae* (8.7%). Overall, the 16S rRNA gene sequencing data corroborated that the bacterial isolates obtained from placental tissues were indeed present on those tissues and were thus not background laboratory contaminants.

**Figure 2.**
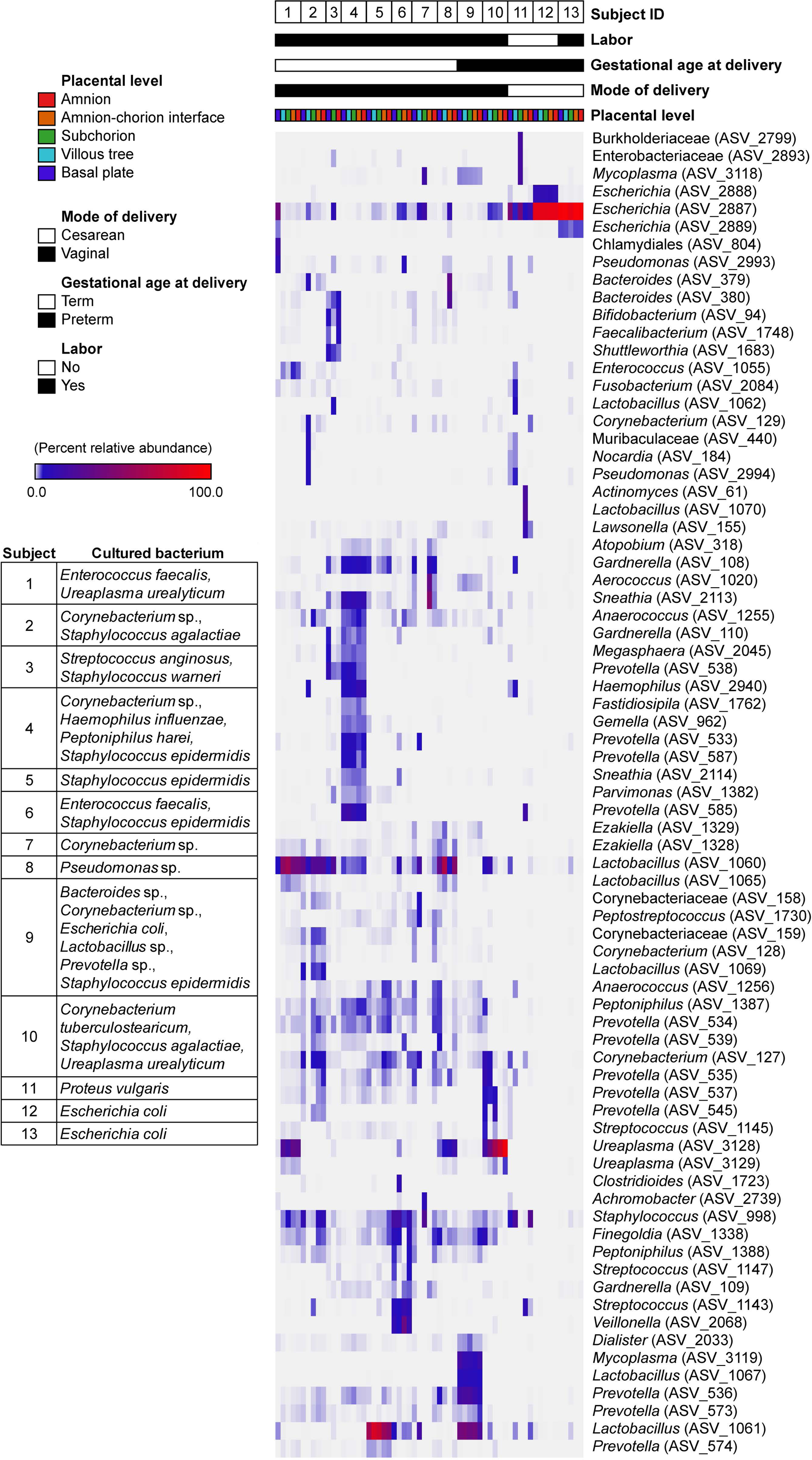
Heat map illustrating concordance between bacterial culture and 16S rRNA gene sequence data from placentas.

### Bacterial taxonomic profiles of placental samples

Investigation of the taxonomic identities of prominent (≥ 2% average relative abundance) ASVs revealed distinct differences in the profiles of placental swabs based on placental level and mode of delivery **(Figure 3 and Figure 4)**. Prior to removing ASVs identified as likely contaminants, *Escherichia* (ASV_2887) was the most relatively abundant taxon in the bacterial profiles of the amnion (5.7%; detected in 60% of samples), subchorion (8.7%; detected in 81% of samples), and villous tree (9.3%; detected in 72% of samples), while *Lactobacillus* (ASV_1061) was most relatively abundant in the bacterial profiles of the amnion-chorion interface (5.9%; detected in 24% of samples) and the basal plate (7.6%; detected in 42% of samples) **(Figure 3)**.

**Figure 3.**
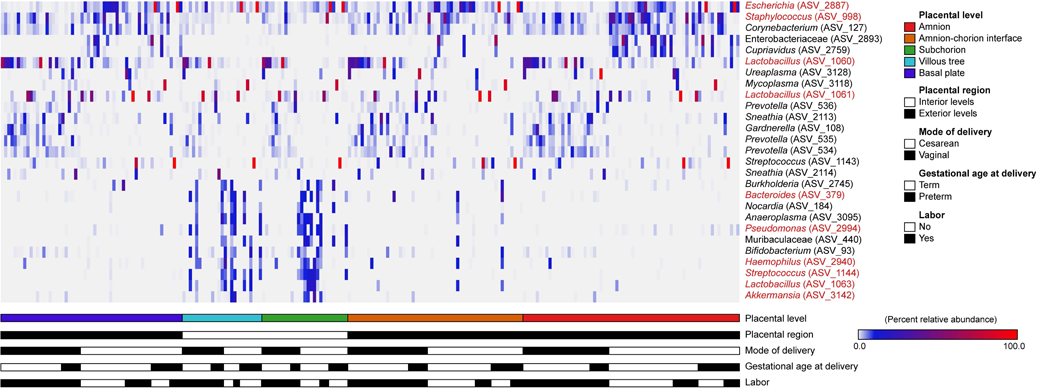
Heat map illustrating the relative abundances of prominent (≥ 2% average relative abundance) amplicon sequence variants (ASVs) among the 16S rRNA gene profiles of placental samples prior to the removal of ASVs identified as likely contaminants. ASVs identified as contaminants are in red font.

**Figure 4.**
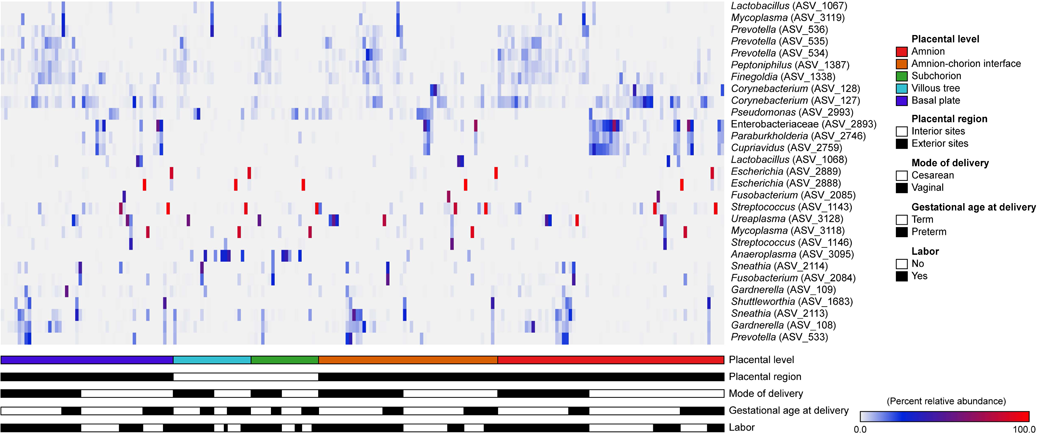
Heat map illustrating the relative abundances of prominent (≥ 2% average relative abundance) amplicon sequence variants (ASVs) among the 16S rRNA gene profiles of placental samples after the removal of ASVs identified as contaminants.

After the ASVs identified as likely background DNA contaminants were removed from the dataset, among vaginally delivered placentas, *Ureaplasma* (ASV_3128) was the most prominent taxon in the bacterial profiles of the amnion (6.4%), amnion-chorion interface (7.2%), subchorion (5.8%), and villous tree (4.8%), while *Sneathia* (ASV_2113) was the most prominent taxon in the bacterial profiles of the basal plate (5.8%) **(Figure 4)**. With respect to distribution (i.e., presence irrespective of relative abundance), five ASVs were widely distributed among all five placental levels: *Anaerococcus* (ASV_1255), *Finegoldia* (ASV_1338), *Gardnerella* (ASV_108), *Peptoniphilus* (ASV_1387), and *Prevotella* (ASV_534) **(Figure 4)**.

Among placentas from cesarean deliveries, *Streptococcus* (ASV_1143) was prominent in the bacterial profiles of all placental levels (> 5%), and *Escherichia* (ASV_2888) was prominent in the amnion-chorion interface (3.6%), subchorion (8.8%), villous tree (8.9%), and basal plate (3.8%) **(Figure 4)**. Two bacterial taxa were only prominent in the profiles of the more exterior sites of the placenta. Specifically, an unclassified Enterobacteriaceae (ASV_2893) was prominent in the amnion (9.5%), amnion-chorion interface (5.2%), and basal plate (5.6%), and *Corynebacterium* (ASV_127) was prominent in the amnion (5.7%) and basal plate (3.5%). *Anaeroplasma* (ASV_3095) was only prominent in the bacterial profiles of the more interior sites of the placenta: subchorion (9.9%) and villous tree (12.4%). With respect to distribution, no ASVs were widely distributed among all five placental levels **(Figure 4)**. Six were widely distributed among the amnion, amnion-chorion interface, and basal plate samples (i.e., exterior sites): Burkholderiaceae (ASV_2746), *Corynebacterium* (ASVs 127, 128, & 129), *Finegoldia* (ASV_1338), and *Pseudomonas* (ASV_2993). Two ASVs were widely distributed only among subchorion and villous tree samples (i.e., interior sites): *Anaeroplasma* (ASV_3095) and Muribaculaceae (ASV_440) **(Figure 4)**.

### Alpha diversity of the bacterial profiles of placental samples

For alpha diversity analyses, ASVs identified as likely background DNA contaminants were removed from the dataset. Linear-mixed-effect modeling **(Table 3)** revealed that the richness (Chao1 index) and the heterogeneity (Shannon diversity) of the bacterial profiles of placental samples differed only by mode of delivery, with vaginally delivered placentas having a higher alpha diversity than cesarean-delivered placentas **(Figure 5)**. When a potential effect of labor was removed from the model addressing bacterial profile heterogeneity (Shannon diversity), there were significant effects of mode of delivery (*X^2^* = 7.62, *df* = 1, *p* = 0.006) and gestational age at delivery (*X^2^* = 4.92, *df* = 1, *p* = 0.027), with placentas from vaginal and term deliveries having higher alpha diversity than those from cesarean and preterm deliveries **(Figure 5)**.

**Figure 5.**
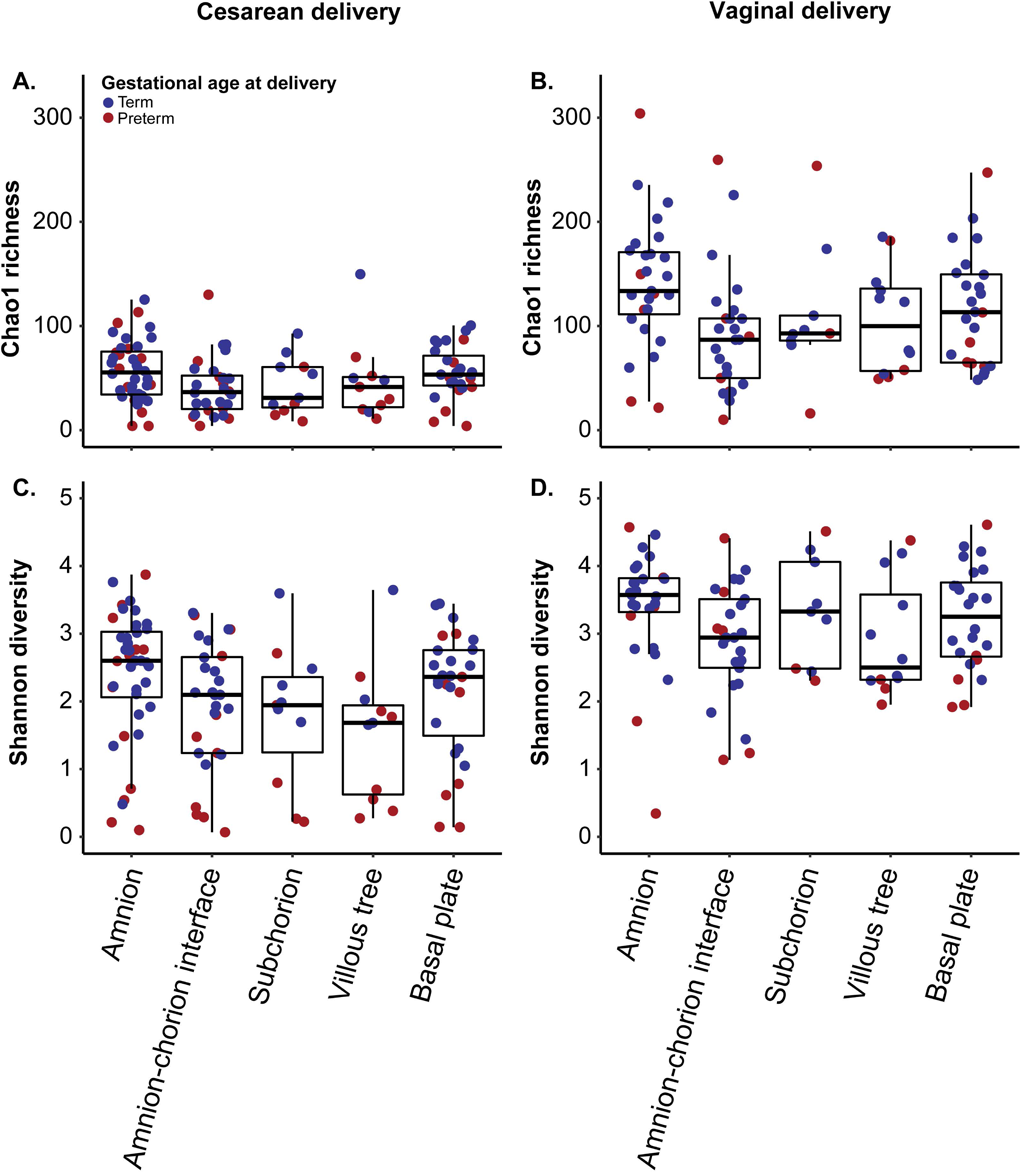
Richness (Chao1 richness) and diversity (Shannon diversity) of placental samples by mode of delivery, gestational age at delivery, and placental level (i.e., amnion, amnion-chorion interface, subchorion, villous tree, basal plate).

**Table 3.** Linear mixed-effect modeling of Chao1 richness and nonparametric Shannon-Wiener diversity of the bacterial profiles of placental samples based on mode of delivery, gestational age at delivery, the presence/absence of labor, and placental level (i.e., amnion, amnion-chorion interface, subchorion, villous tree, and basal plate). Effects were assessed using the models *Richness or diversity = MD*PL + MD*GA + PL*GA + Labor*PL + Labor*GA + Labor*PL + (Patient ID)*, where MD is the mode of delivery, PL is the placental level, and GA is the gestational age at delivery. Patient ID was treated as a random variable.

**Chao1 richness**
| Linear hypothesis | $X^2$ | $df$ | $p$-value |
| --- | --- | --- | --- |
| MD | 8.340 | 1 | **0.004** |
| PL | 3.875 | 4 | 0.423 |
| GA | 0.662 | 1 | 0.416 |
| Labor | 0.036 | 1 | 0.849 |
| MD*PL | 5.820 | 4 | 0.213 |
| MD*GA | 0.026 | 1 | 0.872 |
| PL*GA | 0.744 | 4 | 0.946 |
| PL*Labor | 4.514 | 4 | 0.341 |
| GA*Labor | 0.344 | 1 | 0.558 |

**Shannon diversity**
| Linear hypothesis | $X^2$ | $df$ | $p$-value |
| --- | --- | --- | --- |
| MD | 7.765 | 1 | **0.005** |
| PL | 0.715 | 4 | 0.949 |
| GA | 1.044 | 1 | 0.307 |
| Labor | 0.842 | 1 | 0.359 |
| MD*PL | 2.017 | 4 | 0.733 |
| MD*GA | 0.014 | 1 | 0.905 |
| PL*GA | 0.563 | 4 | 0.967 |
| PL*Labor | 2.178 | 4 | 0.703 |
| GA*Labor | 0.121 | 1 | 0.728 |

### Beta diversity of the bacterial profiles of placental samples

For beta diversity analyses, ASVs identified as likely background DNA contaminants were removed from the dataset. Variation in both the composition (Jaccard index) and structure (Bray-Curtis index) of the bacterial profiles of placental samples was due, in order of influence, to mode of delivery, placental level (i.e., amnion, amnion-chorion interface, subchorion, villous tree, and basal plate), and the presence/absence of labor **(Table 4)**. To address the significant interactions between mode of delivery and placental level as well as mode of delivery and gestational age at delivery, secondary analyses were conducted for each placental level in which the effect of mode of delivery was evaluated separately in term and preterm placentas and any potential effect of gestational age at delivery was assessed separately in placentas from vaginal and cesarean deliveries **(Table 5)**. These analyses revealed that for term placentas, mode of delivery affected the bacterial profiles of each placental level **(Table 5; Figure 6)**. For preterm placentas, wherein sample sizes were smaller and there was thus less power to detect differences, an effect of mode of delivery was evident on the bacterial profiles of the amnion and the basal plate **(Table 5; Figure 6)**. These secondary analyses further revealed that among both vaginally and cesarean-delivered placentas, there was not a consistent effect of gestational age at delivery (i.e., term or preterm) on the overall bacterial profiles of any placental level.

**Figure 6:**
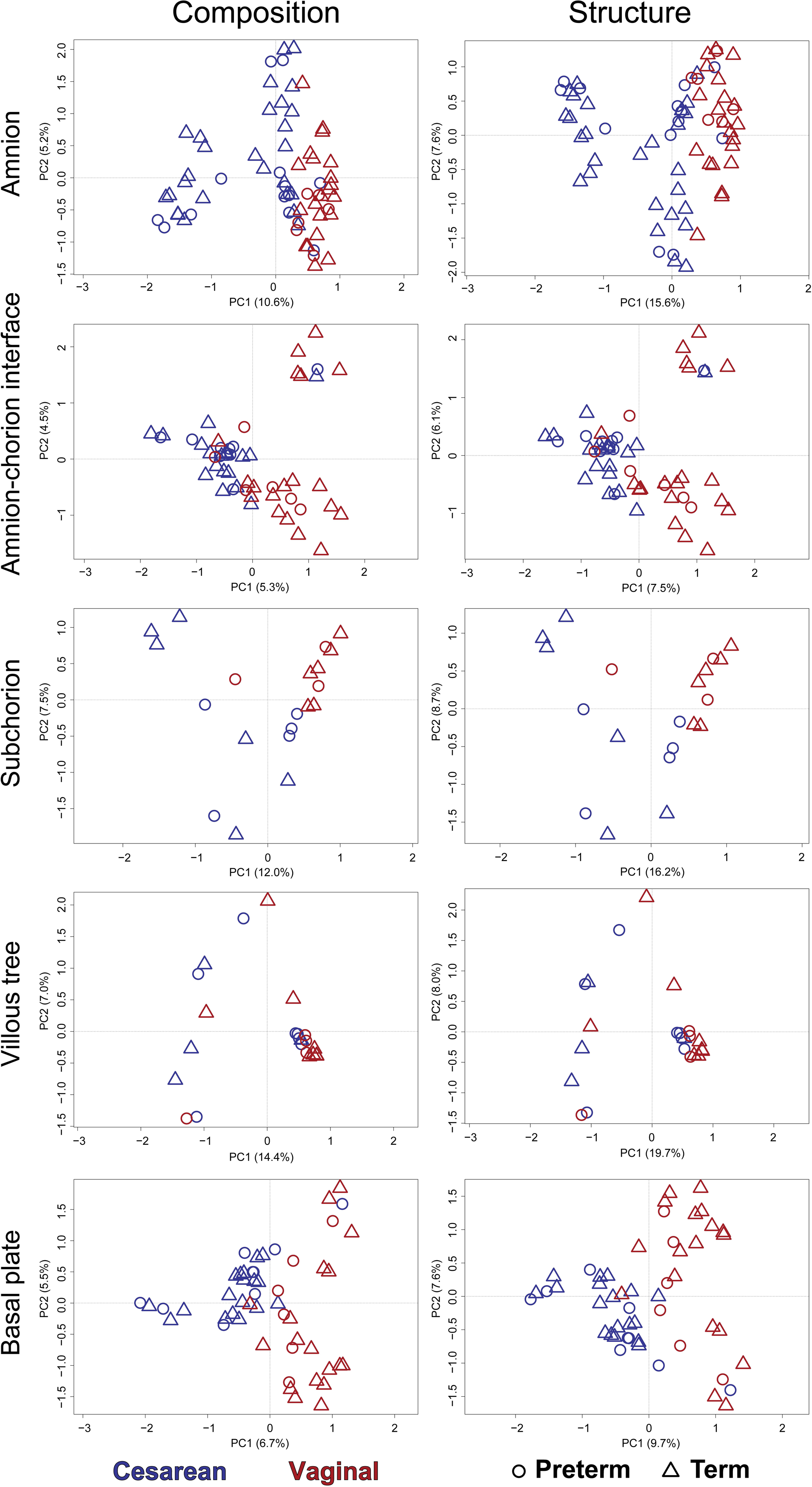
Principal Coordinates Analysis (PCoA) illustrating variation in the composition (Jaccard index) and structure (Bray-Curtis index) of the bacterial profiles of placental samples based on mode of delivery and gestational age at delivery. Given that these analyses were performed for each level of the placenta, there was insufficient power to consider a potential effect of labor.

**Table 4.** PERMANOVA analyses assessing variation in the composition (Jaccard index) and structure (Bray-Curtis index) of the bacterial profiles of placental samples based on mode of delivery, gestational age at delivery, the presence/absence of labor, and placental level (i.e., amnion, amnion-chorion interface, subchorion, villous tree, basal plate). The models were *Composition or structure = MD*PL + MD*GA + PL*GA + Labor*PL + Labor*PL + Labor*GA + (Patient ID)*, where MD is the mode of delivery, GA is the gestational age at delivery, and PL is the placental level. Patient ID was treated as a random variable.

**Composition**
|  | *df* | *F* | *R*<sup>2</sup> | *p*-value |
| --- | --- | --- | --- | --- |
| MD | 1 | 10.447 | 0.046 | **0.001** |
| PL | 4 | 1.35 | 0.024 | **0.001** |
| GA | 1 | 2.419 | 0.011 | 0.324 |
| Labor | 1 | 2.171 | 0.01 | **0.002** |
| MD*PL | 4 | 0.973 | 0.017 | **0.002** |
| MD*GA | 1 | 2.034 | 0.009 | 0.354 |
| PL*GA | 4 | 0.792 | 0.014 | 0.607 |
| PL*Labor | 4 | 0.918 | 0.016 | 0.092 |
| GA*Labor | 1 | 1.554 | 0.007 | 0.341 |

**Structure**
|  | *df* | *F* | *R*<sup>2</sup> | *p*-value |
| --- | --- | --- | --- | --- |
| MD | 1 | 12.847 | 0.055 | **0.001** |
| PL | 4 | 1.764 | 0.03 | **0.001** |
| GA | 1 | 3.024 | 0.013 | 0.918 |
| Labor | 1 | 3.887 | 0.017 | **0.001** |
| MD*PL | 4 | 0.949 | 0.016 | 0.659 |
| MD*GA | 1 | 2.919 | 0.013 | **0.001** |
| PL*GA | 4 | 0.511 | 0.009 | 0.950 |
| PL*Labor | 4 | 0.613 | 0.011 | 0.571 |
| GA*Labor | 1 | 2.259 | 0.01 | 0.738 |

**Table 5.** PERMANOVA analyses assessing variation in the composition (Jaccard index) and structure (Bray-Curtis index) of the bacterial profiles of the placental amnion, amnion-chorion interface, subchorion, villous tree, and basal plate based on mode of delivery and gestational age at delivery.

|  |  | Composition |  |  |  | Structure |  |  |  |
| --- | --- | --- | --- | --- | --- | --- | --- | --- | --- |
|  |  | *df* | *F* | *R*<sup>2</sup> | *p*-value | *df* | *F* | *R*<sup>2</sup> | *p*-value |
| **Vaginal**<br>Term vs.<br>Preterm | Amnion | 26 | 1.115 | 0.043 | 0.191 | 26 | 1.113 | 0.043 | 0.242 |
|  | Amnion-chorion interface | 24 | 1.056 | 0.044 | 0.283 | 24 | 1.063 | 0.044 | 0.334 |
|  | Subchorion | 8 | 1.082 | 0.134 | 0.173 | 8 | 0.993 | 0.124 | 0.464 |
|  | Villous tree | 11 | 0.948 | 0.087 | 0.675 | 11 | 0.931 | 0.085 | 0.603 |
|  | Basal plate | 23 | 1.013 | 0.044 | 0.417 | 23 | 1.100 | 0.048 | 0.278 |
| **Cesarean**<br>Term vs.<br>Preterm | Amnion | 39 | 1.065 | 0.027 | 0.262 | 39 | 0.924 | 0.024 | 0.523 |
|  | Amnion-chorion interface | 27 | 1.043 | 0.039 | 0.320 | 27 | 0.946 | 0.035 | 0.603 |
|  | Subchorion | 10 | 0.095 | 0.095 | 0.776 | 10 | 1.172 | 0.115 | 0.184 |
|  | Villous tree | 10 | 1.173 | 0.115 | 0.109 | 10 | 1.007 | 0.101 | 0.317 |
|  | Basal plate | 26 | 1.102 | 0.042 | 0.146 | 26 | 0.907 | 0.035 | 0.688 |
| **Term**<br>Vaginal<br>vs.<br>Cesarean | Amnion | 47 | 4.714 | 0.093 | **0.001** | 47 | 6.060 | 0.116 | **0.001** |
|  | Amnion-chorion interface | 36 | 2.167 | 0.058 | **0.001** | 36 | 2.328 | 0.062 | **0.001** |
|  | Subchorion | 11 | 1.824 | 0.154 | **0.002** | 11 | 2.507 | 0.200 | **0.003** |
|  | Villous tree | 11 | 1.496 | 0.130 | **0.024** | 11 | 1.811 | 0.153 | **0.043** |
|  | Basal plate | 35 | 3.122 | 0.084 | **0.001** | 35 | 3.379 | 0.090 | **0.001** |
| **Preterm**<br>Vaginal<br>vs.<br>Cesarean | Amnion | 18 | 1.426 | 0.077 | **0.021** | 18 | 1.485 | 0.080 | **0.015** |
|  | Amnion-chorion interface | 15 | 1.001 | 0.067 | 0.451 | 15 | 1.060 | 0.070 | 0.187 |
|  | Subchorion | 7 | 0.953 | 0.137 | 0.632 | 7 | 1.042 | 0.148 | 0.341 |
|  | Villous tree | 10 | 1.035 | 0.103 | 0.334 | 10 | 0.855 | 0.087 | 0.683 |
|  | Basal plate | 14 | 1.419 | 0.098 | **0.008** | 14 | 1.224 | 0.086 | 0.093 |

### Differential abundance of specific taxa in the bacterial profiles of placental samples based on gestational age at delivery

Among placental samples delivered vaginally **(Supplemental Figure 1 and 2)**, both ANCOM-BC and LEfSe analyses identified differentially abundant taxa between term and preterm samples. Overall, most ASVs were enriched in term samples. Specifically, both analyses identified *Gardnerella* (ASVs 108 and 112) as being enriched in the amnion, amnion-chorion interface, and basal plate of term placentas. They also both identified *Corynebacterium* (ASVs 127 and 139) and *Prevotella* (ASVs 537, 549, and 552) as being enriched in the amnion, and *Dialister* (ASV_2032) as being enriched in the villous tree, of term placentas. ANCOM-BC further identified numerous *Lactobacillus* ASVs (1065, 1076, 1077, 1078, 1085, 1087, 1088, 1090, 1096, and 1097) as being enriched in the amnion and amnion-chorion interface of term placentas. BLAST searches (megablast) (69, 70) revealed that all 10 differentially abundant *Lactobacillus* ASVs shared 99.6% nucleotide sequence identity with *Lactobacillus iners* strain DSM 13335 (NR_036982). Very few ASVs were consistently enriched in vaginally delivered preterm placental samples **(Supplemental Figure 1 and 2)**. LEfSe analysis indicated that *Mycoplasma* (ASV_3118) was enriched in the amnion and basal plate of preterm placentas, and *Anaerococcus* (ASV_1265) was enriched in the subchorion and villous tree.

Few differentially abundant taxa were identified among term and preterm placental samples from cesarean deliveries **(Supplemental Figure 3 and 4)**. ANCOM-BC analyses did not identify any bacterial taxa as being enriched in the amnion, amnion-chorion interface, subchorion, or basal plate in term deliveries, and the enrichment of taxa (primarily *Bifidobacterium*, ASV_94) in the villous tree was only very modest. LEfSe analyses indicated that *Pseudomonas* (ASVs 2993 and 2995) was enriched in the amnion, amnion-chorion interface, and basal plate of placentas from term deliveries. No enriched taxa were identified among preterm placental samples from cesarean deliveries in either ANCOM-BC or LEfSe analyses.

### Differential abundance of specific taxa in the bacterial profiles of placental samples based on mode of delivery

Among term placental samples **(Supplemental Figure 5 and 6)**, both ANCOM-BC and LEfSe analyses identified differentially abundant taxa between vaginal and cesarean deliveries. Based on ANCOM-BC analyses **(Supplemental Figure 5)**, *Peptoniphilus* (ASV_1387) and *Prevotella* (ASVs 534-536, 573, 574) were enriched in all levels of term vaginally delivered placentas. *Ezakiella* (ASV_1328) was enriched in all placental levels except for the villous tree, and *Finegoldia* (ASV_1338) was enriched in the amnion, amnion-chorion interface, and the subchorion. *Anaerococcus* (ASVs 1255 and 1256), *Dialister* (ASVs 2031-2033), and *Gardnerella* (ASV_108) were enriched only in the outermost levels (i.e., amnion, amnion-chorion interface, basal plate) of term vaginally delivered placentas. LEfSe analyses also indicated that *Peptoniphilus*, *Prevotella*, *Finegoldia* and *Gardnerella* were enriched in term vaginally delivered placentas, yet they further indicated that *Sneathia* (ASV_2113) was enriched in the amnion, amnion-chorion interface, and basal plate of these placentas **(Supplemental Figure 6)**.

Based on ANCOM-BC analyses **(Supplemental Figure 5)**, no bacterial taxa were enriched in any level of term cesarean-delivered placentas. Based on LEfSe analyses **(Supplemental Figure 6)**, an unclassified Enterobacteriaceae (ASV_2893) was enriched in the outermost levels (i.e., amnion, amnion-chorion interface, basal plate) of term cesarean-delivered placentas. Similarly, *Cupriavidus* (ASV_2759) was enriched in the amnion and basal plate. Conversely, *Anaeroplasma* (ASV_3095) was enriched in the innermost levels (i.e., subchorion, villous tree) of term cesarean-delivered placentas.

Among preterm placental samples **(Supplemental Figure 7 and 8)**, very few bacterial taxa were identified as being differentially abundant between vaginal and cesarean deliveries. Both ANCOM-BC and LEfSe analyses indicated that *Prevotella* (ASVs 534, 536, 539, 603) and *Ureaplasma* (ASV_3128) were enriched in the outermost levels (i.e., amnion and basal plate) of preterm vaginally delivered placentas. ANCOM-BC further identified *Dialister* (ASV_2033), *Finegoldia* (ASV_1338), and *Peptoniphilus* (ASVs 1387 and 1389) as being enriched in the outer levels of these placentas **(Supplemental Figure 7)**.

Based on ANCOM-BC analyses **(Supplemental Figure 7)**, no bacterial taxa were enriched in any level of preterm cesarean-delivered placentas. Based on LEfSe analyses **(Supplemental Figure 8)**, *Mycoplasma* (ASV_3118) was enriched in the amnion of preterm cesarean-delivered placentas.

## DISCUSSION

### Principal findings of the study

1) No bacterial isolates were recovered from placentas from term cesarean deliveries; 2) Placentas from vaginal deliveries were more likely to yield a bacterial isolate than those from cesarean deliveries, and placentas from preterm deliveries were more likely to yield a bacterial isolate than those from term deliveries; 3) Almost all (26/27, 96.3%) bacteria cultured from placental tissues were identified in 16S rRNA gene surveys of those same tissues, indicating they were not laboratory contaminants; 4) The alpha diversity (i.e., heterogeneity) of the 16S rRNA gene profiles of placentas from vaginal and term deliveries was higher than that of placentas from cesarean and preterm deliveries; 5) Variation in the composition and structure of the 16S rRNA gene profiles of placentas was due primarily to mode of delivery and placental level (i.e., amnion, amnion-chorion interface, subchorion, villous tree, basal plate); 6) There was no consistent difference in the overall 16S rRNA gene profiles of any placental level between placentas delivered at term or preterm; 7) Among placentas delivered vaginally, *Corynebacterium*, *Gardnerella*, *Lactobacillus*, and *Prevotella* were enriched in the amnion and/or amnion-chorion interface of those delivered at term; 8) In general, vaginally delivered placentas were consistently enriched in *Finegoldia*, *Gardnerella*, *Peptoniphilus*, and *Prevotella* compared to placentas from cesarean deliveries; 9) Among placentas delivered vaginally, *Mycoplasma* was enriched in the amnion and basal plate, and *Anaerococcus* was enriched in the subchorion and villous tree of those delivered preterm; 10) Among placentas from cesarean deliveries, no bacterial taxa were consistently enriched in either term controls or preterm cases.

### Prior reports of comparisons of the bacterial profiles of term and preterm placentas

Many DNA sequencing-based investigations of placental tissues have suggested an association between specific bacterial DNA signals and preterm birth (1, 9, 10, 48, 49, 52, 53, 56, 71, 72). These associations imply an infectious cause of preterm labor, the potential of which has been reported for multiple bacterial taxa, including *Ureaplasma* and *Mycoplasma* spp., *Fusobacterium nucleatum*, *Gardnerella* spp., *Streptococcus agalactiae*, *Escherichia coli*, *Sneathia* spp., and *Prevotella* spp. (16, 19, 28–34, 37, 73–85). The source of these infectious agents is most commonly vaginal bacteria ascending through the cervix into the intra-amniotic cavity (53, 56, 86–89).

Using DNA sequencing techniques to contrast the bacterial DNA profiles of placental tissues delivered preterm versus at term, multiple studies have concluded that the relative abundances of *Ureaplasma* (9, 48, 52, 53, 71), *Fusobacterium* (48, 52, 53), and *Streptococcus* (9, 48, 52, 56) are positively associated with preterm birth. Yet, besides these three bacterial genera, the specific bacterial taxa identified in placental profiles from preterm deliveries have varied widely across studies, as has the overall structure of the bacterial DNA profiles of preterm placental tissues (7, 10, 48, 49, 52, 71). The reason for these inconsistencies across studies is not clear, however, we propose two potential explanations. First, as noted above, there are many different bacterial taxa which can invade the amniotic cavity and placenta, cause inflammation, and thereby increase the likelihood of preterm birth (48, 74, 87). Indeed, bacterial load (i.e., indicative of colonization) has been associated with inflammation and infection of the fetal membranes in a dose-dependent manner (90). However, the bacteria which invade the placenta are subject specific and thus will be variable within and across studies. Second, even the bacterial profiles of legitimately colonized placental tissues from preterm deliveries are likely to be influenced by underlying methodological differences in general clinical and laboratory environments, sample collection procedures, DNA extraction and sequencing techniques, the bacterial gene or gene region targeted for amplification and sequencing, and the disparate demographics of the different cohorts studied.

These methodological differences among studies also likely explain the large degree of variation in the structure of the bacterial DNA profiles of placental tissues from term deliveries and the bacterial taxa reported to be enriched in term compared to preterm placentas. Specifically, *Escherichia* (1, 72), *Bacteroidetes* (1), *Paenibacillus* (1), *Streptococcus* (10, 48), *Microbacterium*(48), *Rhodococcus* (48), *Corynebacterium*(48), *Acinetobacter* (72), *Enterococcus* (56), *Enterobacter* (52), *Lactobacillus* (52), and *Actinomyces* (10) have all been variably reported as being dominant or enriched in placental tissues from term compared to preterm deliveries. It is intriguing that a bacterial DNA signal can be consistently recovered from placental tissues delivered at term, especially those from cesarean deliveries (1, 4, 10, 50, 54, 55, 57, 58, 72, 91–94), as this suggests the existence of a placental microbiota in human pregnancies in general. However, caution is prudent in interpreting these results. Contemporary bacterial DNA sequencing techniques are highly sensitive, and as such, when working with low microbial biomass samples, such as the placenta, they are inherently susceptible to the influences of background DNA contamination from DNA extraction kits, PCR reagents, and sequencing instruments (3, 6, 7, 9, 11, 59, 60, 95, 96). Therefore, proof of the viability of microorganisms detected in the placenta through DNA sequencing must come from culture and/or transcriptomics of placental tissue for one to conclude the existence of a placental microbiota (11, 65, 97). Indeed, previous studies which included bacterial culture as a complement to DNA sequencing have demonstrated either an absence of bacterial growth (10, 11, 63, 65, 97–99) or bacterial growth that is potentially reflective of delivery- and/or environmental-associated microbiota (2, 24) in placental tissues from term deliveries.

### The findings of this study in the context of prior reports

#### Is there a placental microbiota in term pregnancies?

In this study, through bacterial culture of the human placenta and 16S rRNA gene sequencing of the placental amnion, amnion-chorion interface, subchorion, villous tree, and basal plate, we did not find consistent evidence of a placental microbiota in term pregnancies. Most notably, bacterial isolates were exclusively cultured from placentas from preterm and/or vaginal deliveries – no isolates were obtained from placentas from term cesarean deliveries. This is consistent with recent studies by our group and others indicating that the typical human placenta is unlikely to be inhabited by a viable microbiota (3, 7, 9, 11, 63–65). Notably, here we also demonstrated that the structure of the bacterial DNA profiles of all the sampled levels of placentas (i.e., amnion, amnion-chorion interface, subchorion, villous tree, and basal plate) from term deliveries were significantly affected by mode of delivery. Specifically, term placentas from vaginal deliveries were enriched in *Gardnerella*, *Peptoniphilus*, *Prevotella*, *Anaerococcus*, and *Dialister* compared to placentas from term cesarean deliveries. Each of these bacteria is a common resident of the human vagina (100). These findings reinforce the suggestion that any studies attempting to evaluate the existence of a human placental microbiota must exclusively use placentas obtained from term cesarean deliveries (11). Placentas from term vaginal deliveries, even if the amnion, amnion-chorion, and basal plate are removed prior to bacterial DNA characterization of the subchorion and villous tree, will inherently be contaminated with bacteria and bacterial DNA from the vaginal mucosa.

#### Are there differences in the bacterial profiles of term and preterm placentas?

The absence of a viable placental microbiota in typical human pregnancy does not preclude the importance of bacterial colonization of the placenta for pregnancy complications, especially preterm birth (48, 53, 90). In the current study, one-fourth (5/20) of the cultures of placentas from preterm deliveries yielded bacterial isolates, and all but one of the isolates were further identified in 16S rRNA gene surveys of their respective placentas. This indicates that they were not contaminants introduced in the clinical microbiology laboratory but rather were present on the placenta and viable at the time of sampling. Additionally, four of the five culture-positive placentas delivered preterm yielded cultivable isolates of *Escherichia coli*, *Streptococcus agalactiae*, or *Ureaplasma urealyticum*. *E*. *coli* (90), *Streptococcus agalactiae* (9, 48), and *Ureaplasma urealyticum* (7, 9, 23, 48, 53, 71, 101, 102) have been detected in placentas in prior studies and are established causal agents of preterm birth (86, 103) stillbirth (104), and neonatal sepsis (105) in humans, so it was not surprising that in this study these bacteria were cultured primarily from placentas delivered preterm. Therefore, the results from the culture of placental tissues in this study suggest that viable bacteria were either delivery-associated contamination or that the viable bacteria were likely associated with preterm delivery.

Overall, the bacterial DNA profiles of placental samples were predominated by ASVs which were likely contaminants from DNA extraction kits, PCR reagents, and sequencing instruments (3, 6, 7, 9, 11, 59, 60, 95, 96). After removal of these contaminants, ASVs such as *Mycoplasma* and *Anaerococcus* were enriched in placental tissue from vaginal preterm deliveries, with the former enriching the outer levels (i.e., amnion and basal plate) and the latter enriching the inner levels (i.e., subchorion and villous tree). While there were no specific bacteria consistently enriched in placental tissues from preterm cesarean deliveries, it is important to note that preterm birth is caused, in part, by a suite of pathogenic microorganisms which vary on a case-by-case basis (53, 86). This likely complicates attempts to determine the scope of bacteria enriched in the placenta and amniotic cavity in cases of preterm birth (106).

Notably, however, the alpha diversity of placentas from preterm deliveries was significantly decreased compared to term placentas, suggesting that preterm delivery was associated with the presence of a few bacteria in the placenta. Indeed, decreased richness in placental tissue has been correlated with severe chorioamnionitis and increased bacterial load in placental tissues, both of which are indicative of infection (53). In summary, while a consistent bacterial profile was lacking among placentas from preterm deliveries, the culture and bacterial DNA data from this study were consistent with *E. coli*, *S. agalactiae*, and *Ureaplasma* spp. being variably present in placental tissue, underscoring the varied and complex nature of infectious causes of preterm birth.

### Strengths and limitations of this study

This study has three principal strengths. First, we investigated the bacterial profiles of human placentas simultaneously through culture and 16S rRNA gene sequencing. Second, we incorporated technical controls for potential background DNA contamination in our 16S rRNA gene sequence libraries. Third, we characterized the 16S rRNA gene profiles of five different levels of the human placenta (i.e., amnion, amnion-chorion interface, subchorion, villous tree, basal plate) in the context of term and preterm births. Ultimately, this revealed that vaginal delivery affects the bacterial profiles of all levels of the placenta, not just the amnion, amnion-chorion interface, and the basal plate. Therefore, vaginally delivered placentas should not be used to determine whether there exists a placental microbiota, even if the investigation is limited to the subchorion and villous tree, and investigations of the broader bacterial profiles of placentas from cases of preterm birth should ideally be restricted to those placentas obtained through cesarean delivery.

This study has two primary limitations. First, given that only 62% (214/345) of placental samples from 68 patients yielded a 16S rRNA gene sequence library of at least 500 sequences, we were unable to fully investigate a potential effect of labor on the structure of the bacterial profiles of term and preterm placentas. Further investigation of a potential effect of labor is therefore required. Second, the investigation of placentas was limited to bacterial culture and DNA sequencing without assessment of overall bacterial load or imaging components to demonstrate localization of bacteria potentially associated with preterm birth in different levels of the placenta (9, 11).

### Conclusions

In line with prior investigations, we found no evidence of a microbiota in placentas from term cesarean deliveries. Bacteria were cultured exclusively from placentas from vaginal and/or preterm deliveries. Variation in the structure of placental 16S rRNA gene profiles was due primarily to mode of delivery and the level of the placenta under consideration (i.e., amnion, amnion-chorion interface, subchorion, villous tree, basal plate). The 16S rRNA gene profiles of preterm placentas had less diversity than those of term placentas, suggesting that at least some preterm placentas were populated by potentially infectious bacteria. However, there was not a consistent difference in the composition or structure of the 16S rRNA gene profiles between term and preterm placentas, and among placentas from cesarean deliveries (i.e., those not contaminated by the vaginal delivery process) there were no bacterial taxa consistently enriched in preterm cases. This suggests that the identity of potentially infectious bacteria varied among preterm placentas.

## MATERIALS AND METHODS

### Clinical specimens

Placental samples were obtained at the Perinatology Research Branch, an intramural program of the *Eunice Kennedy Shriver* National Institute of Child Health and Human Development (NICHD), National Institutes of Health, U.S. Department of Health and Human Services, Wayne State University (Detroit, MI), and the Detroit Medical Center (Detroit, MI). The collection and use of human materials for research purposes were approved by the Institutional Review Boards of the NICHD and Wayne State University (#110605MP2F(RCR)). All participating women provided written informed consent prior to sample collection.

### Study design

This was a prospective, cross-sectional, case-control study. The inclusion criteria were: 1) singleton gestation with cesarean or vaginal delivery preterm or at term; 2) patients with vaginal deliveries, cesarean deliveries without labor (NIL), or cesarean deliveries with labor (IL) secondary to obstetrical indications; 3) patients undergoing preterm delivery after an episode of spontaneous preterm labor with intact membranes or preterm prelabor rupture of membranes (PPROM), or secondary to other indications such as preeclampsia or intrauterine growth restriction; and 4) patients providing written informed consent to participate in the study. The exclusion criteria were: 1) any maternal or fetal condition requiring termination of pregnancy; 2) known major fetal anomaly or fetal demise; 3) active vaginal bleeding; 4) multifetal pregnancy; 5) serious medical illness (e.g. renal insufficiency, congestive heart disease, chronic respiratory insufficiency); 6) severe chronic hypertension (requiring medication); 7) asthma requiring systemic steroids; 8) condition requiring anti-platelet or non-steroidal anti-inflammatory drugs; 9) active hepatitis; 10) clinical chorioamnionitis; or 11) antibiotic use within one month of delivery, excluding intraoperative prophylaxis (e.g. Cefazolin for cesarean deliveries and Penicillin for GBS prophylaxis).

### Sample collection

Immediately following delivery, the placenta was placed in a sterile container with a sealed cover and transported to a biological safety cabinet in a nearby research laboratory in Hutzel Women’s Hospital. Therein, swabs (FLOQSwabs, COPAN Diagnostics, Murrieta, CA) were aseptically collected from the amnion, amnion-chorion interface, subchorion, villous tree, and basal plate of the placental disc for molecular microbiology surveys. These swabs were taken from two distinct sites on the placental disc, each site being halfway between the umbilical cord insertion point and the edge of the placental disc. Core placental tissue samples (i.e., ∼1.5 cm^2^ core of tissue from the amnion through to the basal plate) were also collected for bacterial culture. Study personnel wore sterile surgical gowns, hoods, and examination gloves, and used individually packaged, sterile, and disposable scalpels, forceps, and surgical scissors throughout sample collection.

### Bacterial culture of placental tissues and statistical analysis of culture data

Placental tissue samples were transported to the Detroit Medical Center University Laboratories Microbiology Core within anaerobic transport medium surgery packs (Anaerobe Systems, AS-914; Morgan Hill, CA) and 0.85% sterile saline solution tubes (Thermo Scientific, R064448; Waltham, MA) for anaerobic and aerobic bacterial culture, respectively. The placental tissues were processed for culture on the day of collection, as previously described in Theis et al (2019) (11). Briefly, placental tissues were homogenized using a Covidien Precision Disposable Tissue Grinder (3500SA; Minneapolis, MN) and plated via an inoculating loop on three growth media (trypticase soy agar with 5% sheep blood, chocolate agar, MacConkey’s agar) at 35° C under anaerobic (5% CO_2_, 10% H_2_, 85% N_2_) and aerobic (8% CO_2_) atmospheres for four days. A genital mycoplasma cultivation assay (Mycofast US; Logan, UT) was also conducted for each placental tissue sample (107). The taxonomies of resultant isolates were characterized using Matrix-assisted laser desorption/ionization time-of-flight mass spectrometry (MALDI-TOF) (108) within the Detroit Medical Center University Laboratories Microbiology Core.

The effects of mode of delivery (i.e., vaginal/cesarean), gestational age at delivery (i.e., term/preterm), and the presence/absence of labor on the binomial outcome of obtaining a bacterial isolate or not from a placental tissue specimen were assessed through multiple logistic regression using the function for generalized linear models (GLM) in R version 3.6.1 (109). The fit of the logistic models was assessed using the hoslem.test function in the R package ResourceSelection version 0.3.5 (110). Significance of model terms was assessed using the “anova” function (test=“Chisq”). Odds ratios (OR) were then calculated for significant model terms using the epiDisplay package (111).

### DNA extraction from placental swabs

DNA was extracted from placental swabs in a biological safety cabinet using QIAGEN DNeasy PowerLyzer PowerSoil Kits according to the manufacturer’s protocol with minor modifications described in Winters et al (2019) (112). Throughout the DNA extraction process, study personnel wore sterile surgical gowns and examination gloves. To account for potential background DNA contamination, we conducted DNA extractions of 1) six sterile FLOQSwabs processed exactly as the placental swabs and 2) six blank DNA extraction kits that were exposed to the atmosphere of the biological safety cabinet in which DNA extractions took place for 20 minutes. These controls were sequenced alongside placental samples. Ultimately, there were no differences in the 16S rRNA gene profiles of these two types of controls for background DNA contamination (PERMANOVA; Jaccard Index F = 1.035, p = 0.337; Bray-Curtis Index, F = 0.952, p = 0.540), so they were grouped together as “negative controls” for downstream analyses.

### 16S rRNA gene amplification, sequencing, and statistical analysis

Amplification and sequencing of the V4 region of the 16S rRNA gene was performed at the University of Michigan’s Center for Microbial Systems (Ann Arbor, MI) using the dual indexing sequencing strategy developed by Kozich et al (2013) (113). The forward primer was 515F: 5’-GTGCCAGCMGCCGCGGTAA-3’ and the reverse primer was 806R: 5’-GGACTACHVGGGTWTCTAAT-3’. Sequencing libraries were prepared according to Illumina’s protocol for Preparing Libraries for Sequencing on the MiSeq (15039740 Rev. D) and sequencing was conducted using the Illumina MiSeq platform (V2 500 cycles, Illumina MS102-2003) according to the manufacturer’s instructions with modifications found in Kozich et al (2013) (113). Each PCR reaction contained 6.0 µl of template DNA.

Using 35 cycles of standard PCR (for specific PCR conditions, see Theis et al (2019) (11)), 504 of 690 (73%) placental swab samples yielded a 16S rRNA gene amplicon library, as determined by the visualization of PCR products on an E-Gel 96 with SYBR Safe DNA Gel Stain, 2% (Life technologies, Carlsbad, CA). This determination was made by personnel at the University of Michigan’s Center for Microbial Systems, who were blind to the metadata associated with the placental swabs. The remaining 186 placental swabs and the 12 negative controls did not yield usable amplicon libraries at 35 cycles of amplification and were subsequently amplified for 40 cycles and sequenced; 115 placental swabs and 9 negative controls yielded a 16S rRNA gene library of at least 500 sequences. Among these samples amplified for 40 cycles, there was no difference in the composition or structure of 16S rRNA gene profiles between the placental swabs and negative controls from either cesarean (PERMANOVA; Jaccard Index F = 1.134, p = 0.106; Bray-Curtis Index, F = 0.869, p = 0.710) or vaginal deliveries (Jaccard Index F = 1.105, p = 0.142; Bray-Curtis Index, F = 1.013, p = 0.420). Beyond this specific analysis, the placental swabs that only amplified after 40 cycles of PCR were not included in this study, and the data from negative controls were used only to identify potential DNA contaminants in the dataset through conservative use of the *decontam* program (see the section *16S rRNA gene sequence processing and bacterial profile statistical analysis* below) (114).

The binomial outcome of success/failure of 16S rRNA gene amplicon library generation at 35 cycles for placental swabs was assessed using logistic regression with the function for generalized linear mixed-effects models (GLMER) within the lme4 package version 1.1.23 (115) in R version 3.6.1 (109), with the following function options: nAGQ=50 glmerControl(optimizer = “bobyqaℍ, optCtrl = list(maxfun = 100000). Specifically, the likelihood of 16S rRNA gene amplicon library generation from placental swabs was compared among the six patient groups (term cesarean NIL, term cesarean IL, term vaginal, preterm cesarean NIL, preterm cesarean IL, preterm vaginal), as well as between placental level (i.e., amnion, amnion-chorion interface, subchorion, villous tree, and basal plate), with patient ID included in the model as a random variable. Separate models including and excluding labor were evaluated. All possible interaction terms were included in each model. Significance of model terms was assessed by a Type III Wald chi-square test using R package “car” version 3.0.7 (116). A *post hoc* analysis of the effect of placental level was carried out using Tukey’s pairwise comparisons with Bonferroni corrections using the function ‘glht’ in the multcomp package version 1.4.13 (117). Additionally, for testing the binomial outcome of success/failure of 16S rRNA gene amplicon library generation, 16S rRNA gene sequence reads from placental swabs were secondarily grouped by outermost (i.e., amnion, amnion-chorion interface, and basal plate) and innermost (i.e., subchorionic plate and villous tree) regions of the placenta, with the two regions (i.e., outermost and innermost) being characterized by the likelihood of exposure to bacteria in the birth canal and/or delivery room prior to sample processing.

### 16S rRNA gene sequence processing and bacterial profile statistical analysis

16S rRNA gene sequences were clustered into amplicon sequence variants (ASVs), defined by 100% sequence similarity, using DADA2 version 1.12 (118) in R version 3.6.1 (109), according to the online MiSeq protocol (https://benjjneb.github.io/dada2/tutorial.html), with minor modifications as described in Theis et al (2020) (95). The R package *decontam* version 1.6.0 (114) was used to identify ASVs that were likely potential background DNA contaminants based on their distribution among placental swabs and negative controls using the “IsNotContaminant” method. In this study, an ASV was determined to be a contaminant, and was thus removed from the dataset, if it had a *decontam* P score ≥ 0.5 and was present in at least one third of negative controls with an overall average relative abundance of at least 0.5%. Based on these criteria, ten ASVs were identified as DNA contaminants.

Queries of the nucleotide sequences of these 10 ASVs against a curated nucleotide database (rRNA_typestrains/prokaryotic_16S_ribosomal_RNA) using the Basic Local Alignment Search Tool (megablast (69, 70)) returned matches (≥98.8% sequence similarity) for *Bacteroides fragilis* (ASV_379), *Staphylococcus aureus*/*epidermidis* (ASV_998), *Lactobacillus iners* (ASV_1060), *Lactobacillus crispatus*/*gallinarum* (ASV_1061), *Lactobacillus animalis*/*apodemi*/*faecis*/*murinus* (ASV_1063) *Streptococcus pneumoniae*/*pseudopneumoniae* (ASV_1144), *Streptococcus pneumoniae* (ASV_1148), *Haemophilus haemolyticus* (ASV_2940), *Pseudomonas aeruginosa* (ASV_2994), and *Akkermansia muciniphila* (ASV_3142). It is possible that some of these identified contaminants may have actually originated from the placental swabs, as false index barcode pairings can occur during sequence library construction (71, 119, 120). For example, *L. iners* and *L. crispatus* are typical residents of the human vagina (100) and 27/69 (39%) placentas included in this study were obtained following vaginal delivery. Nevertheless, to be conservative, we removed all 10 of these ASVs from the dataset. Additionally, we removed ASV_2887 (*Escherichia*) from the dataset because, although it was not flagged as a contaminant by *decontam*, it was present in all but one control sample and it had an overall average relative abundance of 11%. Prior to the removal of any contaminant ASVs, the dataset contained a total of 10,943,447 sequences and 2,738 ASVs. After the removal of the 11 ASVs deemed to be potential DNA contaminants, 74% of sequences and 2,727 ASVs remained in the dataset.

A preliminary analysis of the 16S rRNA gene sequence data (rarified to 100 sequences per sample) from the replicate swab samples collected from each level of each placenta (i.e., amnion, amnion-chorion interface, subchorion, villous tree, basal plate) revealed that there were significant effects of patient identity and placental level on the bacterial profiles of placentas from both vaginal (PERMANOVA w/ Bray-Curtis index; Patient: F = 9.254, p = 0.001; Level: F = 1.622, p = 0.001) and cesarean (Patient: F = 5.814, p = 0.001; Level: F = 3.238, p = 0.001) deliveries. Indeed, patient identity explained 58.5% and 53.5% of the variation in the 16S rRNA gene profiles of placental samples from vaginal and cesarean deliveries, respectively. To maximize sequence depth and profile coverage, the 16s rRNA gene sequence data from the two replicate swabs collected from each level of each placenta were bioinformatically combined prior to alpha and beta diversity analyses. These combined samples were only included in alpha and beta diversity analyses if they had at least 500 quality-filtered sequences; prior to diversity analyses the combined samples were randomly subsampled to 500 sequences. Ultimately, 214/345 (62.0%) placental samples, from 68 patients, were included in alpha and beta diversity analyses.

Heatmaps of the 16S rRNA gene profiles of placental samples were generated using the open-source software program Morpheus (https://software.broadinstitute.org/morpheus). Alpha diversity of placental samples was characterized using the Chao1 richness and nonparametric Shannon-Wiener diversity (H’; community evenness) indices (121, 122), and variation among patient groups was assessed through linear mixed-effect modeling (LMER) using the lme4 package version 1.1.23 (115) in R version 3.6.1 (109). Alpha diversity indices were calculated in mothur version 1.44.1 (123). Chao1 richness estimates were log-transformed. We considered the predictor variables (mode of delivery, gestational age at delivery (i.e., term/preterm), presence/absence of labor, and placental level) as fixed factors and patient ID as a random variable. All possible interaction terms were included in the models. Residual plots for each model were visually inspected for heteroscedasticity. Significance of model terms was assessed by a Type III Wald chi-square test using the R package “car” version 3.0.7 (116).

Beta diversity of placental samples was characterized using the Jaccard and Bray-Curtis dissimilarity indices. The Jaccard index measures dissimilarities in bacterial profile composition (i.e., presence/absence of each ASV) between samples, and the Bray-Curtis index measures dissimilarities in bacterial profile structure by additionally taking the relative abundance of each ASV into account. Variation in the bacterial profiles of placental samples from different patient groups were visualized through Principal Coordinates Analyses (PCoA) using the R package vegan version 2.5-6 (124). Statistical comparisons of bacterial community composition or structure were made through permutational multivariate analysis of variance (PERMANOVA) (125) using the “adonis” function in the R package vegan version 2.5-6 (124). All possible interaction terms were included in each model and patient ID was controlled for using the “strata” argument.

Variation in the relative abundances of individual ASVs among patient groups was assessed using linear discriminant analysis effect size, or LEfSe (126), with the parameters α = 0.05 and LDA score > 3.5. Prior to LEfSe analyses, singleton and doubleton ASVs were removed from the dataset. Additionally, to identify differentially abundant bacterial ASVs between the profiles of term and preterm placental samples delivered vaginally or via cesarean section, we conducted analysis of composition of microbiomes with bias correction (ANCOM-BC) (127). For ANCOM-BC, placental samples were again only included in analyses if they had at least 500 quality-filtered sequences, however, the dataset was not randomly subsampled to 500 sequences per sample prior to analysis. To correct for multiple comparisons, *p*-values were adjusted using the Benjamini-Hochberg false-discovery rate (FDR). A conservative variance estimate of the test statistic was employed (conserve=TRUE). The level of significance was set to α=0.05.

### Data availability

Sample-specific MiSeq run files have been deposited in the NCBI Sequence Read Archive (PRJNA692425).

## ACKNOWLEDGEMENTS

We thank the physicians and nurses from the Center for Advanced Obstetrical Care and Research and the Intrapartum Unit for their help in collecting human samples. The authors also thank the staff members of PRB Clinical Laboratory, and PRB Histology/Pathology Unit for the processing and examination of placental pathological sections. This research was supported, in part, by the Perinatology Research Branch, Division of Obstetrics and Maternal-Fetal Medicine, Division of Intramural Research, *Eunice Kennedy Shriver* National Institute of Child Health and Human Development, National Institutes of Health, U.S. Department of Health and Human Services (NICHD/NIH/DHHS) under Contract No. HHSN275201300006C. KRT and NG-L were further supported by the Wayne State University Perinatal Research Initiative in Maternal, Perinatal and Child Health. Dr. Romero has contributed to this work as part of his official duties as an employee of the United States Federal Government.

